# Expert Navigators Deploy Rational Complexity-Based Decision Precaching for Large-Scale Real-World Planning

**DOI:** 10.1101/2024.03.25.586612

**Authors:** Pablo Fernandez Velasco, Eva-Maria Griesbauer, Iva Brunec, Jeremy Morley, Ed Manley, Daniel C. McNamee, Hugo J. Spiers

**Author notes:** equal contribution (Pablo Fernandez Velasco, Eva-Maria Griesbauer). equal contribution (Daniel C. McNamee, Hugo J. Spiers).

## Abstract

Efficient planning is a distinctive hallmark of intelligence in humans, who routinely make rapid inferences over complex world contexts. However, studies investigating how humans accomplish this tend to focus on naive participants engaged in simplistic tasks with small state-spaces, which do not reflect the intricacy, ecological validity, and human specialisation in real-world planning. In this study, we examine the street-by-street route planning of London taxi drivers navigating across more than 26,000 streets in London (UK). We explore how planning unfolded dynamically over different phases of journey construction and identify theoretic principles by which these expert human planners rationally precache decisions at prioritised environment states in an early phase of the planning process. In particular, we find that measures of path complexity predict human mental sampling prioritisation dynamics independent of alternative measures derived from the real spatial context being navigated. Our data provide real-world evidence for complexity-driven remote state access within internal models and precaching during human expert route planning in very large structured spaces.

**Significance statement:** Humans can plan efficiently in incredibly complex situations. Existing work has looked at naive participants in simple tasks, which might not be representative of how experts plan in the real world. Here, we study the real-world planning process of London taxi drivers – famous for their expert knowledge of London. By analyzing their response times as a proxy for thinking times, we reveal that at an early stage in their thought process, they store decisions at key street junctions to keep them in mind for later planning. Using computational modeling, we show that taxi drivers prioritize inference at street junctions according to normative metrics measuring how critical a particular decision is for reducing the complexity of planning across the entire city.

## Introduction

In the novel *Invisible Cities*, the author Italo Calvino plays on an analogy between navigating through an urban street network and making moves in a game of chess. Both require the consideration of future actions and their outcomes in order to decide which way to move, that is, planning (1). This analogy between the real and the abstract is a common presupposition in the study of cognitive mechanisms of expert human reasoning (2). Indeed, to date, computational modelling studies of human planning have relied on abstract games such as chess, or Go, or the tower of Hanoi problem for experimental verification (3). Thus, the degree to which theoretic principles developed in the fields of cognitive science and artificial intelligence can be extrapolated to the cognitive computations of human domain experts solving large-scale planning problems in the real world is largely unknown.

The classic approach to planning, both in the cognitive sciences (4–6) and in artificial intelligence (7–9), is to implement a tree search algorithm. This approach involves a representation of possible trajectories across a state-space, with potential actions as branching points. Each sampled trajectory along the decision tree results in an estimated total expected reward or probability of goal achievement, and action plans are decided on based on maximising such estimated quantities (10–12). The fundamental difficulty with algorithms based on tree search is that they become computationally intractable in large state-spaces. This so-called curse of dimensionality (13) presents a puzzle for modelling human cognition, given that humans have the capability to plan in vast state-spaces in which humans operate under substantial time pressures and with limited working memory capacity (14, 15).

And yet, humans routinely defy the curse of dimensionality, by planning successfully in extremely complex environments such as cities (16). Recent approaches in theoretical modelling have sought to specify how humans do this. Existing studies have modelled how people allocate limited computational resources (17, 18), modulate the depth of their planning (6, 19), prune initially unpromising paths (5), and identify how much planning to perform (20). Some of this work has also investigated possible modifications of the problem representation itself, through different forms of simplified representations (21) and task decomposition (22–24) leading to hierarchical planning (14, 25–30).

Efficient planning has been studied in a broad range of experimental and theoretic settings, such as problem-solving games (31, 32), regionalised virtual environments (33, 34) examining the linguistic impact of semantics and preferences for regional crossings (35, 36), latent graphs governing transitions between object stimuli (22, 37, 38) or a virtual subway network task (39). Evidence from these studies suggests not only the existence of regionalised representations of different environments, but also that humans exploit such state-space structure to prioritise particular states in their inference process in order to minimize planning demands (14, 24, 35, 39). However, most studies to date have employed either virtual, small-scale or abstract environments (33–35, 40, 41). As a result, it remains unclear how humans process and exploit state-space structure in large, real-world environments. Cities epitomize the intricacy of planning in real-world environments. If we considered planning a path across a large city that between start and destination involved 30 streets, using a minimal tree-search algorithm would involve an evaluation of over 1 billion potential street sequences (5). The complexity of cities also lies in the fact that they lack definitive regional segregation. For example, in London (UK) there is no definite boundary between *Farringdon* and *City of London*, despite both regions being well known to Londoners (42). In contrast, the state-spaces used in experimental work are often designed so as to accentuate particular state-space structure such as hierarchical decompositions (33–35, 39, 43). A case in point is the study by Balaguer and colleagues (39), in which a virtual subway network was divided into subway lines, which facilitated hierarchically organised planning. Whether the expert human planning in large real-world environments, such as London (Figure 1), without exogenously imposed state-space structure or other cognitive structural representations, remains to be seen.

**FIGURE 1.**
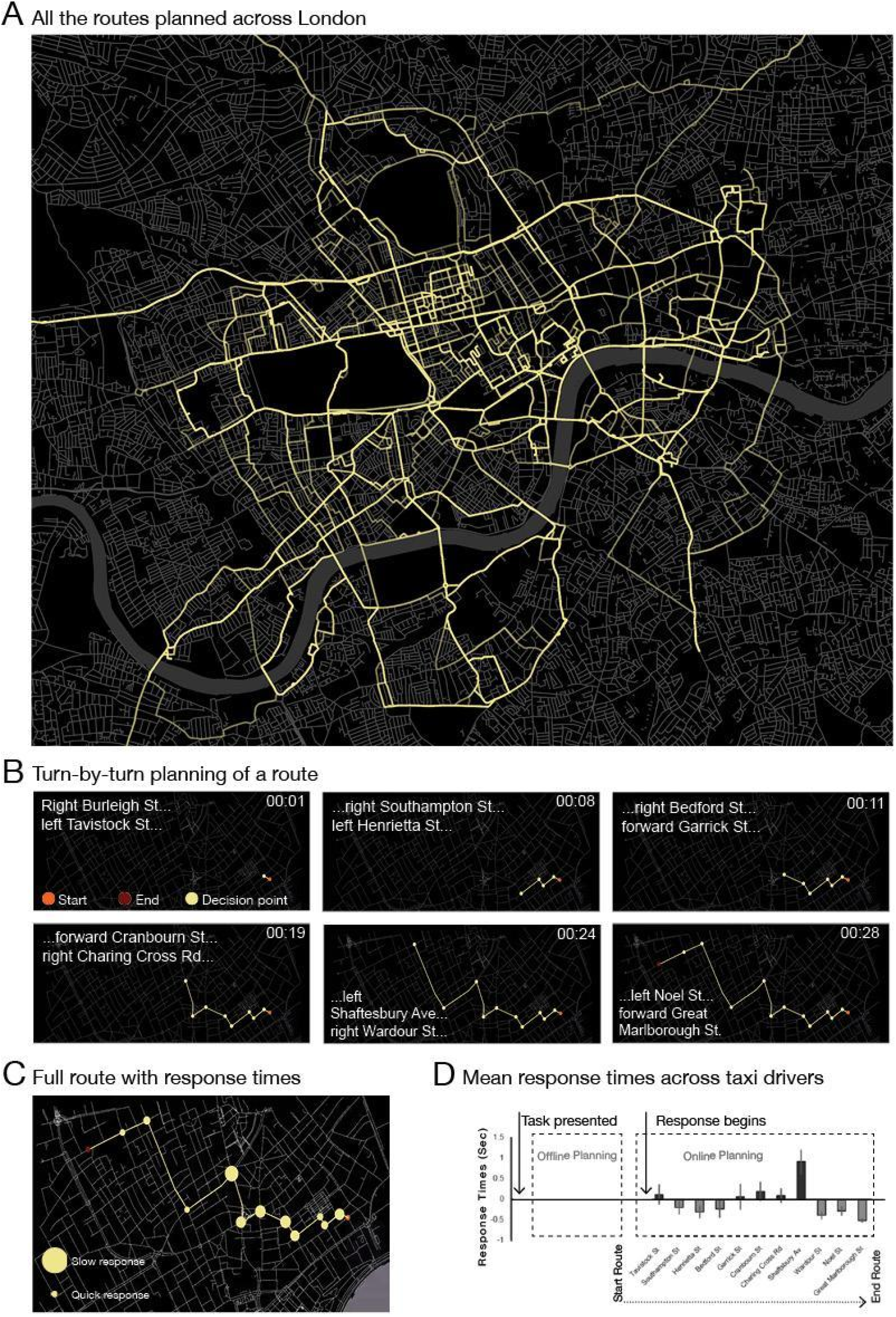
Route planning task. London taxi drivers received pre-recorded audio instructions of the origin and destination locations and called out the names of all the streets they would navigate between the two points, after which they were presented with the next origin-destination pair. All instructions and route plans were audio recorded and transcribed to extract response times. The time lapsed between the instructions and the first recalled street was defined as the offline thinking time (OFF-TT, corresponding to an offline phase of planning). Time lapses between called-out streets of the route were considered as the response times in the online planning phase. **A**) An overlay of the responses from all participants for all routes. For the same origin-destination pair, different participants called different routes. The white rectangle indicates the area covered in route 8 (i.e. as in Fig 1B-1D; task was planning a route from Joe Allen’s restaurant to the Courthouse Hotel). **B)** An example of route 8 (Joe Allen’s restaurant to Courthouse Hotel) for a single participant. The taxi driver “called” the “run” (i.e. route) turn by turn. In the map is the resulting route as it evolved, indicating the corresponding trial times (top-right of each map) and streets called (left). See also Supplementary Movie 1. **C**) The whole route is mapped for the example route shown in B. The circumference size of the circle markers correspond to the length of the time lapses between recalled streets. **D**) The call out times (Response times, average z-score values). Note the long pause before Shaftesbury Avenue that is consistent across all taxi drivers (corresponding to the largest circle in C). See Supplementary Movie 1 for an audio recording of a taxi driver calling out route 8 and an animation of the corresponding route evolving turn-by-turn.

A general quantitative indicator of efficient planning, which may be applied to many such modelling techniques, and may be indexed via the analysis from human reaction times, is the order in which states are processed during planning (44). For example, a simple forward tree search strategy processes states in an ordered fashion directly related to the relative order of those states in the state-space. In contrast, we consider strategies for efficient planning practically result in states being processed out-of-order with respect to the state-space structure. For example, efficient planning with hierarchical representations leverages bottleneck states, which connect contiguous state-space clusters, to be prioritised, and results in a disordered index of states being evaluated with respect to the state-space structure (28). Overall, this approach can reduce the number of states which may need to be evaluated. Our analysis approach is based on identifying evidence for such discontiguous state prioritisation from expert human reaction time data and relating this estimate to information-theoretically motivated variables computed from the state-space graph. In particular, we investigate the influence of two measures of planning complexity on expert human planning which respectively measure how likely and how connected a given state is (14). The first measure is quantified by the successor representation (SR) (45), while the second is measured by the local transition entropy (LTE) in the state-space graph. It can be shown that the product of these two quantities is the contribution of any given state to the *path entropy*. The path entropy is the entropy of the distribution of all paths in the state-space and measures the information-theoretic cost of specifying a particular path e.g. an optimal route to a particular goal. It has been proposed as a measure of planning complexity in large state-spaces and related to neural and behavioural signatures of internal representations for efficient planning (14). Our normative framework proposes that expert humans structure their mental search process by prioritising states in order to maximally decrease path entropy – a principle we refer to as *planning description length minimization*.

Here, we sought real-world evidence of such theoretic principles underlying human expert planning in ecologically valid, large-scale, state-spaces. This effort faces two key challenges, one from an analysis perspective, and the other from an experimental perspective. From an analysis perspective, the size of the state-space presents a significant challenge for fitting and evaluating mechanistic planning models, thus, in this work, we pursued a descriptive statistical approach based on prior normative theoretic analyses of planning efficiency (14). Indeed, this approach was previously used to infer the existence of hierarchical representations in electrophysiological recordings of neural activity and also human sensitivity to critical bottleneck states in laboratory experiments involving the mental simulation of real-world routes (43, 46). A recent successful approach to testing spatial planning has been to examine the reaction times of choices when navigating step-by-step in a fictitious subway network (39). In this approach, increased reaction times imply higher computational demands on the brain to make the next choice and can be employed to infer the computational mechanisms underlying a planning process. Here we extend this approach from a state-space of 35 states to a state space of over 26,000 states: the street network of London (UK). To our knowledge, this is the largest graph structure ever completely enumerated in a human psychological experiment.

From an experimental perspective, the challenge was to identify a large and ecologically valid real-world state-space, and then to recruit a human participant population with expert domain knowledge thereof. To explore how people navigate, ideally participants would tell you their planned route step-by-step. The issue is that the larger the street network, the harder it will be to find participants who can plan routes and recall the names of all the streets between any two points. We propose that licensed London taxi drivers provide an exceptional opportunity to meet this challenge. London taxi drivers are unique in the world for their knowledge of street networks and navigational expertise (16). They are unique with respect to their vast experience and standardised expert training of the London street network (47), and they have been well characterised in prior studies exploring cognition, brain function and structural changes as a result of expertise (48–56). Indeed, in order to obtain their licence, London taxi drivers are specifically examined on their ability to mentally navigate through London by naming all the streets and turns in a planned route between a given origin and destination. This familiar task paradigm forms the basis of our experimental approach (47). Participants were asked to plan routes between a set of origin-destination pairs, and then “call out” the routes step by step. The collected verbal data of the route call-outs was transcribed with regards to street names and to response times. We then employ information-theoretic models to provide predictor variables for the statistical analysis of data collected from forty-three taxi drivers. Specifically, we used general linear models to examine the effect of environmental, cognitive, and theoretical metrics on route planning, with the aim of testing for efficient planning by examining the inferred order in which states are processed. More generally, we assess the impact of different variable classes, such as environmental, spatial, and theoretic, on real-world large-scale expert planning. With response times serving as the behavioural measure, we employed a multivariate linear regression analysis to explore how these variables impacted planning.

## Results

### Data extraction, preparation, and characterisation

Response times were extracted from transcribed verbal reports of London routes (Figure 1, Supplementary Movie 1). Participants were given an origin and a destination, and they had to “call out” the “run” (i.e. the route) step by step (Figure 2A). For each route, the response time data consisted of an *offline thinking time* (OFF-TT, the time from task presentation to the first call-out which characterises the offline phase while the taxi driver silently considers the route) and subsequent *call-out times* (COT, the times between each call-out corresponding to the online phase). We refer to the sum of call-out times for a particular route as the *online thinking time* (ON-TT). Mean offline thinking time was M = 13.83s (*SD* = 13.40) over N = 315 routes. On average, taxi drivers recalled 84.02% of routes. The average total response duration across tasks was M = 17.53s (*SD* = 16.47). The total number of pauses between two consecutively named streets was 3398, with a mean call-out time of M = 1.82s (*SD* = 3.24). For more details of the task see (57). All procedures were approved by the UCL research ethics committee (CPB/2013/150 and EP/2018/008). Informed consent was obtained from all participants.

**FIGURE 2.**
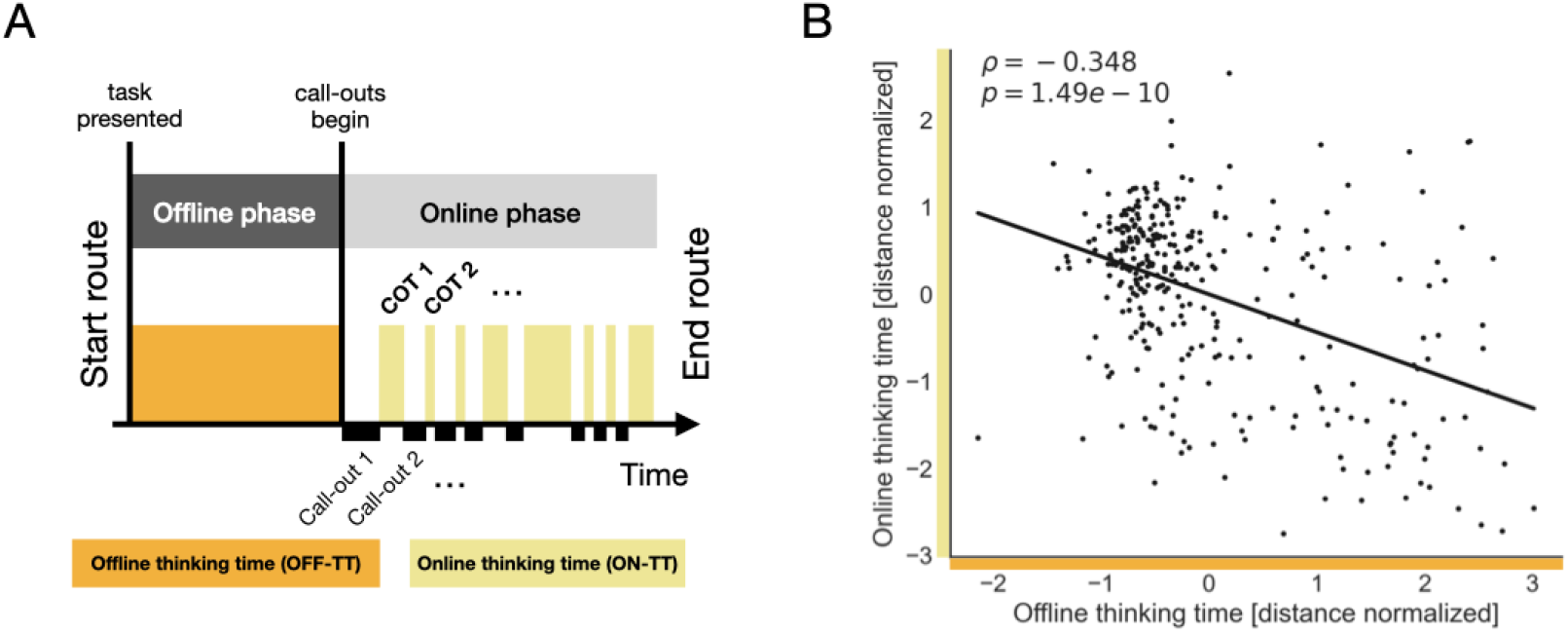
Anti-correlation between offline and online thinking time. **A.** We schematize the trial structure distinguishing the relevant reaction time variables. In particular, we distinguish offline thinking time (OFF-TT) from the call-out times (COTs) in different phases of the trial. The sum of call-out times is referred to as the online thinking time (ON-TT) for the route. **B**. For each run called out by a taxi driver, we normalised the offline thinking time (OFF-TT) and online thinking time (ON-TT) by the streetwise distance of the route. There was a strong negative correlation as measured by Spearman rank correlation *ρ* = -0.348, *p* < 10^−9^. This indicates that longer periods spent thinking offline before the call-outs begin, reduces the burden of planning online. Notably, this correlation was not significant if normalised by the number of steps along each route. This suggests that the physical structure of London was a relevant factor in determining taxi driver planning dynamics.

### Relating thinking times across different route planning phases

In order to examine the relationship between offline OFF-TT and online thinking time ON-TT across routes, we first normalised these data by the streetwise distance on a per-route basis in order to account for the large variability in route length. We then computed the Spearman rank correlation coefficient between OFF-TT and ON-TT (distance normalised) resulting in a strongly significant anti-correlation (, Figure 2B). This indicates that time spent thinking offline reduced the online thinking time required to plan the route. We then tested whether the length of the offline phase differentially impacted online thinking on short versus long timescales. We performed a median-split of our data according to the step number along each route thus defining an EARLY phase of online planning and a LATE online planning phase. There was no significant difference in call-out times in the early versus late online planning phases () and longer offline thinking times did not modulate COTs differently in the early versus late online phase (). This suggests that offline planning operated across the entirety of the route in a global fashion rather than considering streets local to the start position of the route.

### Linear regression analysis of call-out times

We then sought to understand which environmental and theoretic variables influence the modulation of ON-TT via a fixed-effects multivariate linear regression analysis of the call-out times (COTs). This revealed that the following variables had significant effects on call-out times: *Euclidean distance, segment length, remaining streetwise distance-to-destination, remaining direct distance-to-destination, successor representation (SR), segment angular deviation, destination angular deviation*, and *OFF-TT x -SR and OFF-TT x-LTE interactions* (Table 1, OFF-TT is offline thinking time, SR is successor representation, LTE is local transition entropy). See Figure 3A, B for graphical illustrations of the SR and LTE variables. A multivariate correlation analysis was performed which indicated that the theoretic SR and LTE predictor variables were relatively independent of the environmental predictor variables (Figure 3C). A model comparison analysis was performed to compare GLMs expressing distinct hypotheses regarding the interactions between the theoretic variables and taxi driver thinking times (Figure 3D). Table 1 reports the best fitting GLM3 which contains OFF-TT x SR and OFF-TT x LTE interactions. GLM0 included only environmental predictors, GLM1 included environmental and theoretic predictors, and GLMs 2-5 included several distinct combinations of interactions. Notably, the subset of GLMs containing OFF-TT interactions performed much better than the subset of GLMs with OFF-TT interactions thus providing strong evidence for an influence of offline thinking time in determining subsequent call-out times.

**Table 1.** Regression results for general linear model 3 relating offline thinking time to call-out times via interactions between OFF-TT and SR, LTE. The dependent variable is the log-transformed and z-standardised call-out times. Variables that reached significance are highlighted in bold. Consistent with the theoretic analysis of planning complexity based on path entropy, the theoretic variables, namely successor representation (SR) and local transition entropy (LTE), significantly impacted call-out times independent of environmental variables which characterise the veridical spatial context of the London routes.

| Variable | Coef. | Standard Error | t | P>\|t\| | Confidence interval |  |
| --- | --- | --- | --- | --- | --- | --- |
|  |  |  |  |  | [0.025 | 0.975] |
| Intercept | 0.0248 | 0.051 | 0.490 | 0.624 | -0.075 | 0.124 |
| Offline thinking time (OFF-TT) | -0.0197 | 0.023 | -0.858 | 0.391 | -0.065 | 0.025 |
| Plan step number | 0.0025 | 0.001 | 1.752 | 0.080 | -0.000 | 0.005 |
| *Number of planning steps* | -0.0069 | 0.005 | -1.348 | 0.178 | -0.017 | 0.003 |
| *Direct distance-to-destination*<br>*[route total]* | 0.1778 | 0.122 | 1.454 | 0.146 | -0.062 | 0.418 |
| *Direct distance-to-destination*<br>*[remaining]* | -0.1382 | 0.105 | -1.320 | 0.187 | -0.344 | 0.067 |
| ***Streetwise distance-to-destination***<br>***[route total]*** | **-0.5128** | **0.156** | **-3.296** | **0.001** | **-0.818** | **-0.208** |
| ***Streetwise distance-to-destination***<br>***[remaining]*** | **0.3452** | **0.107** | **3.237** | **0.001** | **0.136** | **0.554** |
| ***Street segment length*** | **0.1963** | **0.025** | **7.802** | **0.000** | **0.147** | **0.246** |
| *Segment angular deviation* | 0.0076 | 0.060 | 0.127 | 0.899 | -0.110 | 0.125 |
| ***Destination angular deviation*** | **0.2128** | **0.086** | **2.461** | **0.014** | **0.043** | **0.382** |
| ***Successor representation (SR)*** | **-0.3781** | **0.119** | **-3.186** | **0.001** | **-0.611** | **-0.145** |
| *Local transition entropy (LTE)* | 0.0411 | 0.022 | 1.841 | 0.066 | -0.003 | 0.085 |
| **OFF-TT x SR** | **0.3801** | **0.124** | **3.075** | **0.002** | **0.138** | **0.622** |
| **OFF-TT x LTE** | **-0.0481** | **0.023** | **-2.137** | **0.033** | **-0.092** | **-0.004** |

**FIGURE 3.**
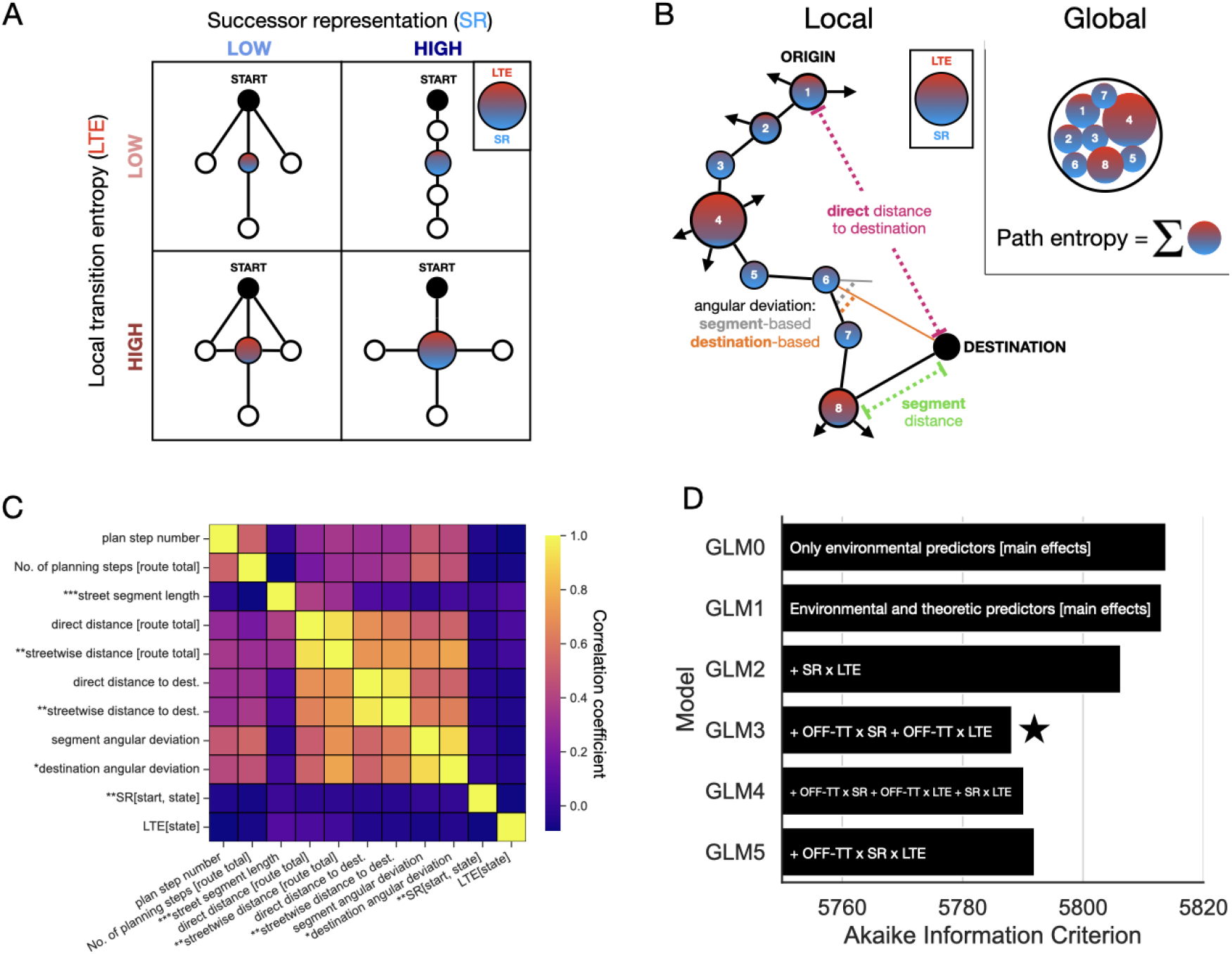
**A.** Graphical depictions of the successor representation (SR) and local transition entropy (LTE) variables. Intuitively, the SR quantitatively captures how common or likely a particular state is while LTE reflects the complexity of a decision at an environmental state. **B**. Environmental and theoretic predictor variables are schematized along an origin-destination route from local and global perspectives. SR and LTE jointly specify the local decision complexity along a run. Path entropy is the global sum of these local complexity measurements. **C**. This predictor variable correlation matrix demonstrates that the theoretic predictors, the successor representation (SR) and local transition entropy (LTE), varied independently with respect to each other and the environmental predictor variables. **D**. Using the Akaike information criterion (AIC) for model comparison (a lower AIC value indicates a better explanatory model), the best explanatory model of the taxi driver call-out times was GLM3 (starred) containing main effects of the environmental and theoretic predictors as well as interaction terms between OFF-TT and the SR and LTE variables. Using the Bayesian information criterion instead of AIC led to the same result. This indicates that the SR and LTE variables influence state prioritisation during the offline planning phase before the taxi driver starts calling out the route.

Since the combination of SR and LTE determine a normative localised metric of planning complexity (Figure 3B), we review in detail how they impacted call-out times (14). There was a highly significant and negative main effect of SR on call-out times such that more predictable states were associated with faster COTs. This effect did not interact with the planning step number (p=0.16) indicating that it was consistently present through the planning process rather than being related to e.g. cognitive fatigue. The main effect of LTE trended strongly towards slowing COTs but did not reach significance (p=0.066).

### Multi-phase dynamics in response times consistent with efficient discontiguous state access during planning

In our experiment, planning can take place before the first call-out (the offline phase) and then, may continue in parallel while the route is verbally reported (Figure 2A). We sought to examine how SR and LTE influence behavioural response times from a dynamic multi-phase perspective over the course of planning. LTE measures the local processing cost at a particular street segment, and it is weighted by SR in determining its contribution to global planning complexity according to the planning description length perspective (see Methods, Eqn. 1). In a state-space as complex as London, there are inevitably too many states to consider and search through assuming limited availability of memory capacity and time. Given this, our theoretic analysis prescribes a rational strategy whereby high SR and high LTE states are prioritised for planning (Figure 4C).

**FIGURE 4.**
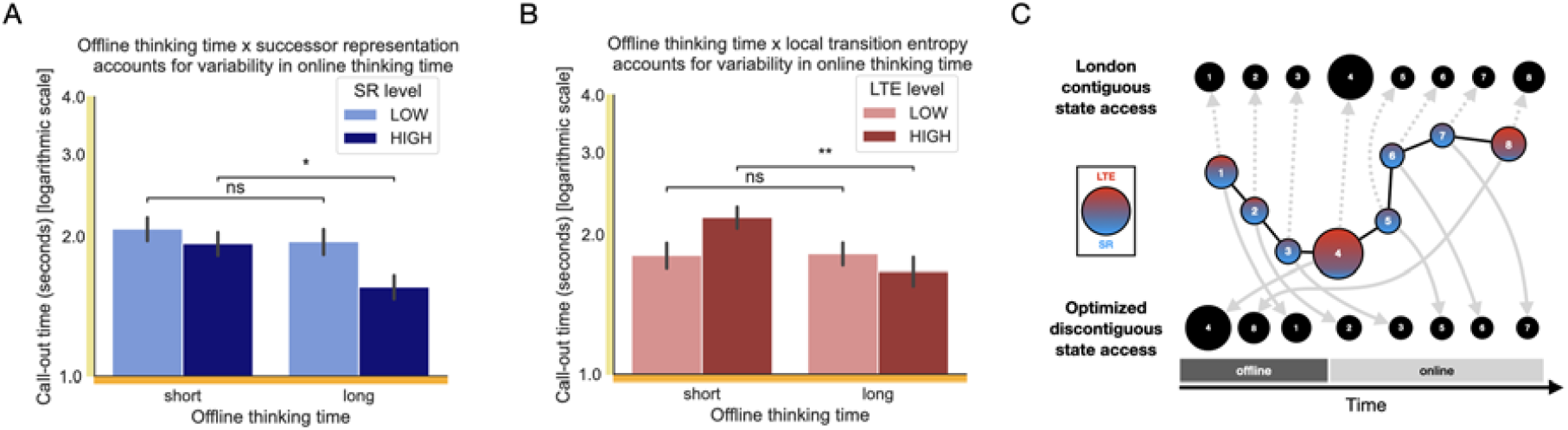
The local complexity of the street network, measured by successor representation and local transition entropy, interacts with offline thinking time in predicting variability in taxi driver call-out times. **A.** Call-out times specifically for highly predictive states were faster if the taxi driver spent more time thinking offline prior to beginning to verbally report the route (two-sided t-test between median splits, p=0.02, t=2.565). **B**. Furthermore, the offline thinking time interacted with the theoretic LTE variable, measuring local decision complexity, in a similar manner (two-sided t-test between median splits, p=0.005, t=3.045). Specifically, a longer period of offline thinking resulted in significantly decreased online COT for complex local transition structures (high LTE). **C**. Our data points to a spatially discontiguous global prioritisation of remote states as planning evolves whereby high SR and high LTE states are accessed expeditiously offline and the associated local decision precached (solid-line arrows). This stands in contrast to an ordered sequential structure to London-contiguous state sampling in simulation-based models of planning such as tree search (dashed-line arrows).

In order to test this, we focused on the relationship between the length of the offline planning phase and the subsequent effects of SR and LTE on call-out times. We reasoned that a longer offline planning phase should be optimised to prospectively plan specifically at high SR and high LTE states and precache the associated decisions leading to a reduction in the subsequent call-out times at these states (Figure 4A, B). Indeed, our model comparison (Figure 3D) identified significant interactions between the amount of time the taxi drivers deployed to think offline and the SR and LTE variables in determining subsequent call-out times. Furthermore, this hypothesis was directly tested using median-splits of the data according to the length of the offline thinking period and comparing high and low SR (LTE) call-outs across this split. For longer offline planning periods, we found a significant and selective reduction in call-out time for both the SR (Figure 4A) and LTE (Figure 4B) variables. In contrast, a planning process based on sequential state roll-outs e.g. tree search, whereby states are prioritised in the order they occur along the route, would predict an interaction effect of planning step number and the length of the offline planning period (Figure 4C). In particular, the call-out times would be relatively reduced for states closer to the origin; however such an effect was not present in the data (p=0.648, t(43)=-0.457). Importantly, we verified that there were no robust autocorrelations in the SR (1-lag autocorrelation = 0.0) and LTE (1-lag autocorrelation = 0.1) variables along the evaluated routes in order to rule out a confound whereby the SR and LTE variables were structured in a spatially consistent manner with respect to the London state-space.

## Discussion

In the present study, we set out to examine expert planning in a large-scale, ecologically valid, setting by testing licensed taxi drivers engaged in navigation tasks across London. London taxi drivers are uniquely positioned for this investigation because they undergo rigorous training and have a vast, first-hand experience of the street network. In our study, we asked forty-three taxi drivers to mentally plan routes between a set of origin-destination pairs while verbally describing those routes street by street. We then transcribed their responses and measured the response times before and while calling out the names of streets and turns along routes through the city. Using a multivariate linear regression analysis, we examined how a range of spatial, environmental, and theoretic variables impacted planning with response times serving as the behavioural measure. This approach exposed three principles of efficient expert planning, namely predictive mapping, complexity-based prioritisation, and global non-sequential decision precaching.

We sought to determine how planning dynamically unfolded over different phases of each trial. Specifically, by examining the influence of the time length of the offline planning period on call-out times, we aimed to identify how expert planners rationally organise their planning process in time by selectively targeting, potentially remote, environment states for decision precaching. This involves retrieving the relevant state information for prioritised offline planning, and storing the resulting local plan for that state, thus enabling faster plan retrieval during the online planning phase. In order to minimise planning description length, states should be accessed according to the SR and LTE measures as knowledge of the correct transitions at such states provide the maximal reduction in the overall uncertainty regarding a particular route (14, 58). In accordance with this idea, the data we collected from expert London taxi drivers reflected selective reductions in online response times at high SR and high LTE states for longer offline planning periods. The SR and LTE measures in our theory (14) are respectively analogous to the so-called need and gain terms described in theoretic work based on the Q-DYNA variant of model-based RL based on sequential state rollouts for planning (59). In this model, the gain term refers to the amount of reward that can be gained by implementing a Bellman backup update to the policy. This requires that a particular reward function be specified (60). Given that our task, and the real-world experience of taxi drivers, corresponds to the multi-task RL setting with different starts and goals defining a huge number of possible reward functions, we suggest that the LTE measure provides an appropriate analogous measure in the multi-task setting agnostic to any particular reward function. Indeed, the conceptual usage of LTE in our model is consistent with a number of information-theoretic studies investigating efficient generic state-space representations for exploration and planning in artificial agents (22, 61–64). An important distinction with respect to meta-optimised models based on Q-DYNA and tree search (6, 59), is the apparent global non-sequential, spatially discontiguous, nature of the prioritisation effect we observe. In our data, the effect of route step number did not reach significance and the associated coefficient was very weak. Therefore, states were apparently not accessed in the order they could be sequentially generated from the origin but out-of-order with respect depending on their SR and LTE values. We suggest that such a global re-organization of planning from local simulations to non-sequential decision precaching for critical states throughout the London state-space may reflect extremely refined planning strategies for expert planners. This predicts alternative forms of neural replay which may be testable in data and motivates the investigation of possible generative models of planning which exhibit such memory access dynamics (65, 66).

High LTE states tend to form critical junction states which interface between adjacent clusters and thus the relatively slow call-outs at these states may be caused by internal switches between distinct local state-space representations (14). An illustrative example is the turn at Shaftesbury Avenue in Figure 1, a critical junction state that corresponds to a very slow reaction time. This is consistent with our theoretical perspective given that the LTE for the Shaftesbury Avenue state in the London state-space is in the 95th percentile of the LTE distribution across the entire city. Thus, expert navigators, such as taxi drivers, may exploit segmentations of London into hierarchically organised regions in order to optimise for planning demands (35, 39). This result provides real-world support to previous theories of hierarchical planning coming from both computational and laboratory work (14, 26–30).

In our analysis, we included a range of environmental variables focusing on those highlighted in studies on human psychology in urban environments (33, 67). Some of these variables, such as distances-to-destination, may reflect a navigation strategy based on “counting down” distance to a target location as observed in homing animals and insects, after apparently approximately path-integrating outbound trajectories (68). Notably, only distance-to-destination metrics measured along the traversed streets modulated the taxi drivers’ call-out times. Direct distances-to-destination, which ignore the street structure, did not. This suggests that taxi drivers have internally metrized London based on their natural behavioural affordances, i.e. driving through the streets, rather than absolute Euclidean distance. The theoretic predictor variables in our model were selected based on the mathematical definition of planning description length. However, this does not rule out the possibility that variables derived from other planning models may explain some of the variance in the taxi driver thinking times or provide alternative interpretations. For example, a taxi driver entering a very familiar region of London may use a regional representation with SR values conditioned on the entrance state to this region providing a better individualised fit to his or her thinking. In general, we expect this unique experimental population of human experts to provide further opportunities to test a range of models.

Another environmental measure that has a significant effect on response times is segment length, i.e. the length of the current segment at the state-level. This effect means that taxi drivers are slower at transitioning through physically longer segments. A potential explanation for this effect is that taxi drivers employ some form of mental visualisation of streets along the route based on an internal model of the physical properties of London (58, 69). This is consistent with existing work suggesting that taxi drivers imagine in-street views of the trajectory when computing route plans (47). Further evidence supporting the hypothesis that taxi drivers planned using internal representations which reflected the embodied spatial structure of London comes from the fact that destination angular deviation (i.e. angular deviation with respect to the final destination) also has an effect on response times. We can see this metric as a measure of the curvature of the route within a real-world spatial embedding as opposed to a pure graph-based representation of the state-space which lacks angular relationships. Within this framework, the slowing down of response times as a function of this variable is consistent with vector-based navigation (70). Using deep reinforcement learning, Banino and colleagues provided support for the idea that entorhinal grid cells contribute the spatial metric enabling the calculation of vectors between the current location and the goal (71). Another recent computational approach employed a hybrid model that combined a map-based strategy reliant on boundary cells and place cells and a vector-based strategy reliant on grid-cells (72, 73). Future work could investigate how the use of vector-based navigation, predictive maps, non-sequential precaching, and hierarchical planning vary from task to task, from environment to environment, and from participant to participant. Indeed, regarding the latter, factors which presumably influence inter-individuality in the behaviour and planning processes of taxi drivers are route preferences, regional familiarity, and prior distributions of routes navigated. In future work, we aim to investigate such variability in human expert planning using large-scale databases of taxi driver route distributions and demand indicators.

## Methods

### Experimental protocol

Taxi drivers were presented with an origin-destination pair and then were required to verbally describe a route from the origin to the destination within London. These verbal descriptions of the routes were recorded and transcribed from the taxi drivers (57). Response times were calculated as pauses between two consecutively named streets (see Figure 1 highlighted in yellow). The response time to the first named street, which tended to be significantly longer, is referred to as the *offline thinking time (OFF-TT)* while the subsequent response times are referred to as *call-out times (COT)*. The former is conceptualised as characterising the *offline* planning phase before the driver begins announcing the route whereas the latter corresponds to the *online* planning phase while the driver verbally navigates the route. The sum of call-out times for a particular route is referred to as the *online thinking time (ON-TT)*. Response times and thus were log-transformed prior to linear regression analyses and presentation. Furthermore, these data were standardised via z-scoring on a per-participant basis.

Origin-destination combinations were selected such that the corresponding routes varied in their environmental and structural properties (e.g. Euclidean distance, path length, direction of travel, and planning complexity). The experimental data was collected in two phases. In the first phase, tasks consisted of 12 origin-destination pairs. In the second data collection phase, tasks consisted of 8 origin-destination pairs, including two origin-destination pairs from the original set to allow for a comparison across the two groups. To keep traffic conditions consistent, participants were asked to imagine they were carrying out the planning task at 11.00 AM on a typical Monday. The data from both experiments was treated as one data set, because the Wilcoxon Signed-rank test indicated no group differences between the two sets of taxi drivers for tasks 7 and 8 for log-transformed (Mdn(S1) = -0.22, IQR = 1.18, Mdn(S2) = -0.16, IQR = 1.24, p = .252, r = .046) or z-standardised (Mdn(S1) = -0.29, IQR = 1.11, Mdn(S2) = -0.33, IQR = 1.24, p = .688, r = .016) call-out times.

### Environmental measures of London spatial context

To test the impact of spatial features of the street network on route planning behaviour, *call-out times* (COTs) were analysed in association with the following environmental metrics: direct *distance-to-destination, streetwise distance-to-destination, segment length, remaining direct distance-to-destination, remaining streetwise distance-to-destination, segment angular deviation, destination angular deviation. Direct distance-to-destination* corresponds to the Euclidean distance from origin to destination as the crow flies, at the route level. *Streetwise distance-to-destination* corresponds to the distance from origin to destination along the shortest path, at the route level. *Segment length* corresponds to the length of the current segment, at the state-level. *Remaining streetwise (direct) distance-to-destination* corresponds to the remaining streetwise (direct) distance to the destination. *Segment angular deviation* corresponds to the angular deviation with respect to the next segment in a route. *Destination angular deviation* corresponds to the angular deviation with respect to the destination of the route. See Figure 3B for a schematic illustration of these environmental measures.

### Planning description length: a normative measure of planning complexity

We use path entropy as an information-theoretic measure of planning complexity under a stochastic search model (14). This quantifies how much information must be generated on average in order to specify a route from an origin state to a destination state. The matrix *H*(*τ*|*o*) of path entropy for all possible trajectories *τ* emanating from an origin state *o* in London and can be expressed as a matrix-vector product of the *successor representation (SR) M* and the *local transition entropy* (LTE) *H* (74, 75):

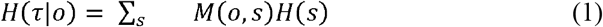

where *s* is an arbitrary London state. The *successor representation (SR)* is defined as:

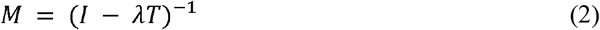

where *T* is the transition matrix defining the probability of transitioning between states of the environment under a random walk (i.e. exploratory) policy. Specifically, *T*(*s, s*′) is the probability moving to state *s*′ from state *s*. The SR measures how likely different states (in our case, street segments) are to be sampled under a given transition policy and a model of the environment (45). To a state s, we associate the SR measure *M*(*o, s*) where *o* is the origin state for a particular route task. Note that the SR definition (Equation 2) does not depend on a particular goal state in contrast to previous work where a goal-absorbing SR was computed for every goal state (14). Given that any location in London may be a goal state for a taxi driver then this goal-agnostic SR serves as a close approximation but obviates the need to compute a distinct SR for every goal state. *Local transition entropy (LTE)* is defined as

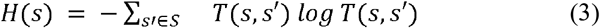

In our analyses, the SR and LTE values associated with a reported London position are computed with respect to the transition structure at the previously reported position. That is, for a transition *s* → *s*′ then the LTE H(s) is aligned with the call-out time for *s*′ as this is expected to reflect the complexity of the planning process which produced state *s*′ (see Figure 1). Since the overall complexity of planning may be expressed as a product of SR and LTE, we reasoned that these variables should determine the mental labour of planning and, consequently, guide how planning is efficiently structured in expert planners under resource limitations (76–78). From a normative perspective, we suggest that plans are processed at states in the order by which they reduce the overall uncertainty in the route trajectory (14). We refer to this as a *planning description length minimization* strategy. For example, taxi drivers may establish an approximate transition plan for a critical junction such as Charing Cross before establishing the sub-route from the origin to Charing Cross.

### Theoretic variables and state-space network structure

The SR and LTE variables have graphical interpretation in terms of the structure of the state-space (Figure 3A) (14, 79). For example, from the graph-theoretic perspective, high SR-LTE states tend to form critical junction states (Figure 3A, bottom right) while high SR low-LTE correspond to highly connected that may be concatenated into sub-routes (Figure 3A, top right) as in the options framework in hierarchical reinforcement learning (RL) (28, 80). The network structure interpretation of SR and LTE is related to their role as normative metrics of planning complexity. Indeed, the correspondence between the network structure motifs described by these theoretic variables and their normative computational implications has been studied previously (22, 64) and their interactions explain call-out time variability (Sup. Figure 1A). Taken together, these complementary perspectives suggest resource-rational principles by which planning may be re-organized from a localised serial search approach to a globalised multi-scale process guided by SR and LTE metrics in extremely large state-spaces (Figure 4C).

### Statistical analysis

To test the effects of variables from the street network on route planning, we employed multivariate linear regression models assuming fixed effects across taxi drivers. The dependent variable was the log-transformed response times from the route call-outs (COTs) which was z-scored on a per-driver basis. We explored thirteen independent variables. These included the spatial metrics outlined above: *Euclidean distance-to-destination, streetwise distance-to-destination, segment length, remaining streetwise distance-to-destination, segment angular deviation, destination angular deviation*. The independent variables in the models also included the following: *offline thinking time (OFF-TT), plan step number, number of planning steps, successor representation measurement* (SR), *local transition entropy (LTE)*, and *SR-LTE interaction. Plan step number* corresponded to the callout number within a route, at the state-level. *Number of planning steps* corresponded to the total number of callouts in a route, at the route-level.

### Model comparison

We fitted several general linear models (GLM) to predict taxi driver call-out times (COTs). Each GLM had a qualitatively distinct set of predictor variables examining a specific relationship between the offline thinking time (OFF-TT) and the theoretic predictors SR and LTE in predicting COT. GLM0 contained only spatial predictor variables while GLM1 contained both spatial and theoretically predicted variables (see Figure 3C). GLMs 2-5 contained interaction terms involving the theoretic SR and LTE variables and OFF-TT. The best fitting model included interaction terms between OFF-TT and each of the SR and LTE theoretic variables. The nature of these significant interactions is further investigated in Figure 4. Notably, this model comparison serves to provide strong evidence of interactions between the SR and LTE variables and offline thinking time OFF-TT given that the class of models containing interactions (GLMs 3-5) provides a much stronger fit than the model class without (GLMs 0-2).

## Data, Materials, and Software Availability

Data and the analysis code can be found on: https://github.com/dmcnamee/taxi-driver-planning

## ACKNOWLEDGEMENTS

PFV’s work was supported by the British Academy (PFSS23\230053). DM’s work was supported by the Champalimaud Foundation. We thank Francesco Trapani, Margarida Sousa, and the anonymous reviewers for their feedback.

## Notes

https://github.com/dmcnamee/taxi-driver-planning

### Competing Interest Statement

The authors have declared no competing interest.

### Summary of Updates

We realised that the term "precaching" is better suited to describe the results than "prioritization" (the term employed in the previous version).

## References

1. I. Calvino, Invisible cities (Houghton Mifflin Harcourt, 1978).

2. A. Newell, H. A. Simon, Human problem solving (Prentice-Hall, 1972).

3. K. R. Allen, et al., Using Games to Understand the Mind. (2024). 10.31234/osf.io/hbsvj.

4. N. D. Daw, Y. Niv, P. Dayan, Uncertainty-based competition between prefrontal and dorsolateral striatal systems for behavioral control. Nat. Neurosci. 8, 1704–1711 (2005).

5. Q. J. M. Huys, et al., Bonsai Trees in Your Head: How the Pavlovian System Sculpts Goal-Directed Choices by Pruning Decision Trees. PLOS Comput. Biol. 8, e1002410 (2012).

6. B. van Opheusden, et al., Expertise increases planning depth in human gameplay. Nature 618, 1000–1005 (2023).

7. C. E. Shannon, Programming a computer for playing chess. Lond. Edinb. Dublin Philos. Mag. J. Sci. 41, 256–275 (1950).

8. D. Silver, et al., Mastering the game of Go with deep neural networks and tree search. Nature 529, 484–489 (2016).

9. S. Russell, P. Norvig, Artificial Intelligence: a modern approach, 4th US ed. Univ. Calif. Berkeley (2021).

10. L. A. Streeter, D. Vitello, A Profile of Drivers’ Map-Reading Abilities. Hum. Factors 28, 223–239 (1986).

11. R. J. Elliott, M. E. Lesk, Route finding in street maps by computers and people in Proceedings of the Second AAAI Conference on Artificial Intelligence, AAAI’82., (AAAI Press, 1982), pp. 258– 261.

12. K. J. Miller, S. J. C. Venditto, Multi-step planning in the brain. Curr. Opin. Behav. Sci. 38, 29–39 (2021).

13. R. Bellman, Dynamic programming (Princeton University Press, 1957).

14. D. McNamee, D. M. Wolpert, M. Lengyel, Efficient state-space modularization for planning: theory, behavioral and neural signatures in Advances in Neural Information Processing Systems, (Curran Associates, Inc., 2016).

15. T. L. Griffiths, Understanding Human Intelligence through Human Limitations. Trends Cogn. Sci. 24, 873–883 (2020).

16. P. Fernandez Velasco, H. J. Spiers, Wayfinding across ocean and tundra: what traditional cultures teach us about navigation. Trends Cogn. Sci. 28, 56–71 (2024).

17. J. Snider, D. Lee, H. Poizner, S. Gepshtein, Prospective Optimization with Limited Resources. PLOS Comput. Biol. 11, e1004501 (2015).

18. F. Callaway, et al., A resource-rational analysis of human planning in Proceedings of the 40th Annual Meeting of the Cognitive Science Society, CogSci 2018, (The Cognitive Science Society, 2018), pp. 178–183.

19. M. Keramati, P. Smittenaar, R. J. Dolan, P. Dayan, Adaptive integration of habits into depth-limited planning defines a habitual-goal–directed spectrum. Proc. Natl. Acad. Sci. 113, 12868– 12873 (2016).

20. E. Russek, D. Acosta-Kane, B. van Opheusden, M. G. Mattar, T. Griffiths, Time spent thinking in online chess reflects the value of computation. (2022). 10.31234/osf.io/8j9zx.

21. M. K. Ho, et al., People construct simplified mental representations to plan. Nature 606, 129–136 (2022).

22. A. Solway, et al., Optimal Behavioral Hierarchy. PLOS Comput. Biol. 10, e1003779 (2014).

23. Q. J. M. Huys, et al., Interplay of approximate planning strategies. Proc. Natl. Acad. Sci. 112, 3098–3103 (2015).

24. C. G. Correa, M. K. Ho, F. Callaway, N. D. Daw, T. L. Griffiths, Humans decompose tasks by trading off utility and computational cost. PLOS Comput. Biol. 19, e1011087 (2023).

25. K. S. Lashley, “The problem of serial order in behavior” in Cerebral Mechanisms in Behavior; the Hixon Symposium, (Wiley, 1951), pp. 112–146.

26. M. M. Botvinick, Hierarchical reinforcement learning and decision making. Curr. Opin. Neurobiol. 22, 956–962 (2012).

27. J. P. O’Doherty, S. W. Lee, D. McNamee, The structure of reinforcement-learning mechanisms in the human brain. Curr. Opin. Behav. Sci. 1, 94–100 (2015).

28. M. M. Botvinick, Y. Niv, A. G. Barto, Hierarchically organized behavior and its neural foundations: A reinforcement learning perspective. Cognition 113, 262–280 (2009).

29. M. S. Tomov, S. Yagati, A. Kumar, W. Yang, S. J. Gershman, Discovery of hierarchical representations for efficient planning. PLOS Comput. Biol. 16, e1007594 (2020).

30. H. Bast, et al., “Route Planning in Transportation Networks” in Algorithm Engineering: Selected Results and Surveys, Lecture Notes in Computer Science., L. Kliemann, P. Sanders, Eds. (Springer International Publishing, 2016), pp. 19–80.

31. G. Ward, A. Allport, Planning and Problem solving Using the Five disc Tower of London Task. Q. J. Exp. Psychol. Sect. A 50, 49–78 (1997).

32. C. A. Knoblock, “A Theory of Abstraction for Hierarchical Planning” in Change of Representation and Inductive Bias, The Kluwer International Series in Engineering and Computer Science., D. P. Benjamin, Ed. (Springer US, 1990), pp. 81–104.

33. J. M. Wiener, H. A. Mallot, “Fine-to-Coarse” Route Planning and Navigation in Regionalized Environments. Spat. Cogn. Comput. 3, 331–358 (2003).

34. J. M. Wiener, A. Schnee, H. A. Mallot, Use and interaction of navigation strategies in regionalized environments. J. Environ. Psychol. 24, 475–493 (2004).

35. W. Schick, M. Halfmann, G. Hardiess, F. Hamm, H. A. Mallot, Language cues in the formation of hierarchical representations of space. Spat. Cogn. Comput. 19, 252–281 (2019).

36. W. Schick, Acquisition and consolidation of hierarchical representations of space. (2018). 10.15496/publikation-23395.

37. J. M. Wiener, S. J. Büchner, C. Hölscher, Taxonomy of human wayfinding tasks: A knowledge-based approach. Spat. Cogn. Comput. 9, 152–165 (2009).

38. B. Hommel, J. Gehrke, L. Knuf, Hierarchical coding in the perception and memory of spatial layouts. Psychol. Res. 64, 1–10 (2000).

39. J. Balaguer, H. Spiers, D. Hassabis, C. Summerfield, Neural Mechanisms of Hierarchical Planning in a Virtual Subway Network. Neuron 90, 893–903 (2016).

40. S. Büchner, et al., Path choice heuristics for navigation related to mental representations of a building in Proceedings of the European Cognitive Science Conference, (Taylor and Francis Delphi, Greece, 2007), pp. 504–509.

41. D. Badre, A. S. Kayser, M. D’Esposito, Frontal Cortex and the Discovery of Abstract Action Rules. Neuron 66, 315–326 (2010).

42. E.-M. Griesbauer, E. Manley, D. McNamee, J. Morley, H. Spiers, What determines a boundary for navigating a complex street network: evidence from London taxi drivers. J. Navig. 75, 15–34 (2022).

43. A. E. G. F. Arnold, G. Iaria, A. D. Ekstrom, Mental simulation of routes during navigation involves adaptive temporal compression. Cognition 157, 14–23 (2016).

44. M. G. Mattar, M. Lengyel, Planning in the brain. Neuron 110, 914–934 (2022).

45. P. Dayan, Improving Generalization for Temporal Difference Learning: The Successor Representation. Neural Comput. 5, 613–624 (1993).

46. K. Bonasia, J. Blommesteyn, M. Moscovitch, Memory and navigation: Compression of space varies with route length and turns. Hippocampus 26, 9–12 (2016).

47. E.-M. Griesbauer, E. Manley, J. M. Wiener, H. J. Spiers, London taxi drivers: A review of neurocognitive studies and an exploration of how they build their cognitive map of London. Hippocampus 32, 3–20 (2022).

48. E. A. Maguire, et al., Navigation-related structural change in the hippocampi of taxi drivers. Proc. Natl. Acad. Sci. 97, 4398–4403 (2000).

49. E. A. Maguire, K. Woollett, H. J. Spiers, London taxi drivers and bus drivers: A structural MRI and neuropsychological analysis. Hippocampus 16, 1091–1101 (2006).

50. E. A. Maguire, R. Nannery, H. J. Spiers, Navigation around London by a taxi driver with bilateral hippocampal lesions. Brain 129, 2894–2907 (2006).

51. H. J. Spiers, E. A. Maguire, Spontaneous mentalizing during an interactive real world task: an fMRI study. Neuropsychologia 44, 1674–1682 (2006).

52. H. J. Spiers, E. A. Maguire, Thoughts, behaviour, and brain dynamics during navigation in the real world. Neuroimage 31, 1826–1840 (2006).

53. H. J. Spiers, E. A. Maguire, The neuroscience of remote spatial memory: a tale of two cities. Neuroscience 149, 7–27 (2007).

54. H. J. Spiers, E. A. Maguire, Neural substrates of driving behaviour. Neuroimage 36, 245–255 (2007).

55. H. J. Spiers, E. A. Maguire, A navigational guidance system in the human brain. Hippocampus 17, 618–626 (2007).

56. H. J. Spiers, E. A. Maguire, The dynamic nature of cognition during wayfinding. J. Environ. Psychol. 28, 232–249 (2008).

57. E.-M. Griesbauer, et al., London taxi drivers exploit neighbourhood boundaries for hierarchical route planning. bioRxiv 2024–02 (2024).

58. D. McNamee, D. M. Wolpert, Internal Models in Biological Control. Annu. Rev. Control Robot. Auton. Syst. 2, 339–364 (2019).

59. M. G. Mattar, N. D. Daw, Prioritized memory access explains planning and hippocampal replay. Nat. Neurosci. 21, 1609–1617 (2018).

60. R. S. Sutton, Dyna, an integrated architecture for learning, planning, and reacting. ACM SIGART Bull. 2, 160–163 (1991).

61. C. Salge, C. Glackin, D. Polani, Empowerment --an Introduction. [Preprint] (2013). Available at: http://arxiv.org/abs/1310.1863 [Accessed 4 December 2023].

62. K. Gregor, D. J. Rezende, D. Wierstra, Variational Intrinsic Control. [Preprint] (2016). Available at: http://arxiv.org/abs/1611.07507 [Accessed 4 December 2023].

63. A. Goyal, et al., InfoBot: Transfer and Exploration via the Information Bottleneck. [Preprint] (2023). Available at: http://arxiv.org/abs/1901.10902 [Accessed 22 March 2024].

64. K. Archer, N. C. Volpi, F. Bröker, D. Polani, A space of goals: the cognitive geometry of informationally bounded agents. [Preprint] (2022). Available at: http://arxiv.org/abs/2111.03699 [Accessed 4 December 2023].

65. A. E. Comrie, et al., Hippocampal representations of alternative possibilities are flexibly generated to meet cognitive demands. bioRxiv 2024.09.23.613567 (2024). 10.1101/2024.09.23.613567.

66. D. C. McNamee, The generative neural microdynamics of cognitive processing. Curr. Opin. Neurobiol. 85, 102855 (2024).

67. D. Yesiltepe, et al., Entropy and a sub-group of geometric measures of paths predict the navigability of an environment. Cognition 236, 105443 (2023).

68. M. Müller, R. Wehner, Path integration in desert ants, Cataglyphis fortis. Proc. Natl. Acad. Sci. 85, 5287–5290 (1988).

69. P. W. Battaglia, J. B. Hamrick, J. B. Tenenbaum, Simulation as an engine of physical scene understanding. Proc. Natl. Acad. Sci. 110, 18327–18332 (2013).

70. F. L. Moël, T. Stone, M. Lihoreau, A. Wystrach, B. Webb, The Central Complex as a Potential Substrate for Vector Based Navigation. Front. Psychol. 10 (2019).

71. A. Banino, et al., Vector-based navigation using grid-like representations in artificial agents. Nature 557, 429–433 (2018).

72. V. Edvardsen, A. Bicanski, N. Burgess, Navigating with grid and place cells in cluttered environments. Hippocampus 30, 220–232 (2020).

73. N. Nyberg, É. Duvelle, C. Barry, H. J. Spiers, Spatial goal coding in the hippocampal formation. Neuron 110, 394–422 (2022).

74. L. Ekroot, T. M. Cover, The entropy of Markov trajectories. IEEE Trans. Inf. Theory 39, 1418– 1421 (1993).

75. M. Kafsi, M. Grossglauser, P. Thiran, The Entropy of Conditional Markov Trajectories. IEEE Trans. Inf. Theory 59, 5577–5583 (2013).

76. S. J. Gershman, E. J. Horvitz, J. B. Tenenbaum, Computational rationality: A converging paradigm for intelligence in brains, minds, and machines. Science 349, 273–278 (2015).

77. W. Kool, M. Botvinick, Mental labour. Nat. Hum. Behav. 2, 899–908 (2018).

78. F. Lieder, T. L. Griffiths, Resource-rational analysis: Understanding human cognition as the optimal use of limited computational resources. Behav. Brain Sci. 43, e1 (2020).

79. K. L. Stachenfeld, M. M. Botvinick, S. J. Gershman, The hippocampus as a predictive map. Nat. Neurosci. 20, 1643–1653 (2017).

80. Y. Jinnai, D. Abel, D. Hershkowitz, M. Littman, G. Konidaris, Finding Options that Minimize Planning Time in Proceedings of the 36th International Conference on Machine Learning, (PMLR, 2019), pp. 3120–3129.

