## Supplementary Movie 1 for "Expert Navigators Deploy Rational Complexity-Based Decision Precaching for Large-Scale Real-World Planning"

### Supplementary Material

#### Supplementary Figures

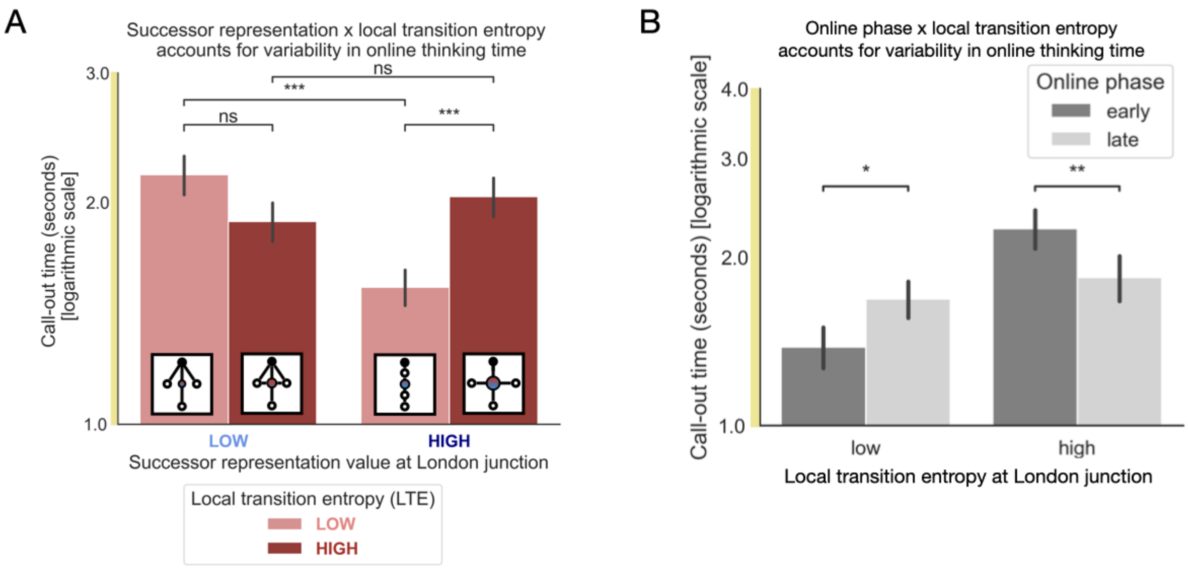


**SUPPLEMENTARY FIGURE 1. A.** In GLM4, there was a significant interaction between SR and LTE in predicting taxi driver call-out times (p=0.003). This effect was driven by a reduction in COT specifically for low LTE street transitions with high SR measurements. That is, streets with relatively uncomplicated local structure and high throughput under random walk dynamics. Successive street segments of this type are normatively chunked together under hierarchical control models. One interpretation of this interaction is that the taxi drivers prioritise the chunking of more common multi-street sequences in memory as opposed to all multi-street sequences due to memory resource limitations. Error bars reflect standard error of the mean. **B.** We also analysed the progression of planning through different phases of the online planning period. In particular, COTs specifically for high complexity transitions become faster as planning progresses (p=0.006). Furthermore, the offline planning time (OPT) differentially interacts with theoretic variables (SR and LTE) in determining call-out times (COTs) between planning phases. In particular, an interaction between OFF-TT and LTE (p=0.0001) and a three-way interaction between planning step number, OFF-TT, and LTE (p=0.015) were observed. A longer offline planning period resulted in a significantly decreased COT specifically for complex local transition structures (high LTE). This effect was particularly pronounced in the early planning phase consistent with the continuous evolution of planning throughout the initial, early, and late planning phases. Error bars reflect standard error of the mean.
